# Cell-Scale Biophysical Cues from Collagen Fiber Architecture Instruct Cell Behavior and the Propagation of Mechanosensory Signals

**DOI:** 10.1101/2020.08.12.248179

**Authors:** Joseph M. Szulczewski, David R. Inman, Maria Proestaki, Jacob Notbohm, Brian M. Burkel, Suzanne M. Ponik

## Abstract

Mechanosensory cues from the extracellular matrix underpin numerous cellular behaviors including tumor cell migration yet are influenced by the local structure and organization of the matrix in unknown ways. To investigate mechanical cues with respect to local collagen organization, we used a combination of intravital imaging of the mammary tumor microenvironment and 3D collagen gel systems with both migratory MDA-MB-231 cells and acellular pNIPAAm beads. We identified that fiber organization directs a bias in cell response along the axis of alignment. Using innovative methodology, we determined that local collagen alignment resulted in a 30-fold difference in directional cell-scale stiffness and also dramatically altered the rate at which cell-induced fiber displacements decayed over distance. Our results reveal differential mechanical properties across orthogonal directions in aligned matrices that provide sizeable cues to the cell and have important implications for cellular mechanosensing and cell-cell communication within the tissue microenvironment.

## Introduction

Cells use physical features of the microenvironment to calibrate their gene expression and morphology (*1–3*), and to organize their internal machinery to produce forces for biological processes like cell migration, which occurs in development, homeostasis, and disease. During these processes, complex and multifactorial cues from the matrix provide feedback to the cells that determine the distribution of cellular contractile components and modulate the magnitude of the cell response. While our understanding of cellular mechanosensing during migration typically focuses on the role of focal adhesions and the molecular clutch (*4–6*), the role of the surrounding matrix in limiting or promoting future cellular behavior remains less clear. Little is known about how the organization of cell machinery and cell-applied forces deform their environment, or how these deformations may provide necessary feedback for the greater mechanosensory system.

Physiological environments like those of the native stroma or tumor microenvironment have a fibrous structure and, importantly, the architecture of the fibers is known to impact disease progression. In breast cancer, alignment of collagen fibers perpendicular to the tumor boundary is prognostic of worse patient outcome (*7*). A factor underlying the relationship between fiber alignment and disease progression is an increase in persistent cell migration (*8*), and aligned fibers have been shown to promote persistent migration by limiting the distribution and shape of focal adhesions (*9*). To better understand the relationship between cellular behavior and matrix architecture, it is necessary to examine the mechanical properties of the fibers in various organizations. As cells contract the matrix, the fibers under tension become more aligned while those under compression buckle (*10*). This creates local anisotropy in stiffness, and explains why fibers that are aligned parallel to the cell axis are thought to be stiffer than their perpendicular counterparts (*11*). The anisotropic property of fibers provides valuable feedback to the cell, but because the mechanics of fibrous matrices are complicated and evolve over time, even the magnitude of stiffness between perpendicular and parallel fiber orientations is unknown.

The fibrous structure and organization of the ECM may also affect how the matrix deforms due to the forces produced by the cell. This has implications for mechanical communication that occurs between cells. Recently, macrophages were observed to respond to the pulsatile contractions of myofibroblasts during wound healing (*12*). The directed migration of macrophages was not dependent on matrix stiffness, but rather on matrix deformations produced by a myofibroblast. This is significant because in contrast with previous studies that only considered the cell response to the matrix, this observation is an example where matrix response to the cell is equally important. Given the heterogeneity in fiber organization within physiological tissues, the question remains whether the waves of fiber deformation propagate uniformly throughout the matrix.

Both static cues (i.e. stiffness) provided by the structure of the ECM and more dynamic cues (i.e. matrix deformations) resulting from cellular contractions are important for efficiently organizing cellular machinery used during numerous cellular processes. In this study, we characterized how the different matrix organizations respond to localized cell-scale forces and influence migratory behavior. Using intravital imaging of transgenic and xenograft murine mammary carcinoma models, we established that as cancer cells migrate through their local microenvironment they simultaneously deform both individual fibers and the surrounding fiber network. We then developed an *in vitro* model system to reproducibly introduce local, strain-induced alignment within a 3D collagen gel. With this model, we controlled migratory behavior and quantified the relative magnitude of durotactic cues in both random and aligned matrices. We also clearly demonstrated how fiber organization affects the propagation of displacements, which could have implications for cellular mechanosensing and cell-cell communication within the matrix. Through this study, we aimed to generate a more comprehensive perspective of the cellular mechanosensing system. This includes both the physical cues that direct and orient cellular adhesions to efficiently organize and apply forces, and the characteristics of the subsequent cell-scale matrix deformations that may instruct cellular behavior. Together, this work highlights the importance of local structure and organization in understanding cellular mechanosensing within physiological, 3D environments.

## Results

### Migrating cancer cells physically deform and reorganize collagen fiber architecture in vivo

To determine the nature and extent of how mechanical cues condition future cellular behavior in a physiological environment, we built upon an intravital imaging system to visualize cells interacting with native ECM of a MMTV-PyMT (PyMT) murine mammary tumor model (*13*). Using this system, collagen fibers were visualized through second harmonic generation (SHG) while endogenous cellular fluorescence was used to track cell movement in the tumor. Initial observations of early stage tumors highlighted the vast heterogeneity of the fiber network within the tumor microenvironment. Within a single tumor, different fields of view within close proximity to each other often exhibited vastly different fiber configurations. In regions of early tumor initiation, collagen fibers had a relaxed appearance with curly, random fibers distributed throughout the field of view (Fig 1a). Typically, both tumor (*) and adipose (+) cells could be easily identified. Near larger tumor masses or between separate foci of the growing mass, fibers appeared straightened and under tension (Fig 1a). Local regions of aligned fibers could be identified; propagating either parallel or perpendicular to the tumor boundary (Fig 1a, arrows). Importantly, this observation was consistent with previous studies from human samples where the orientation of aligned fibers perpendicular to the tumor boundary correlated with poor disease-specific and disease-free survival in patients with invasive ductal carcinoma (*7*).

**Figure 1:**
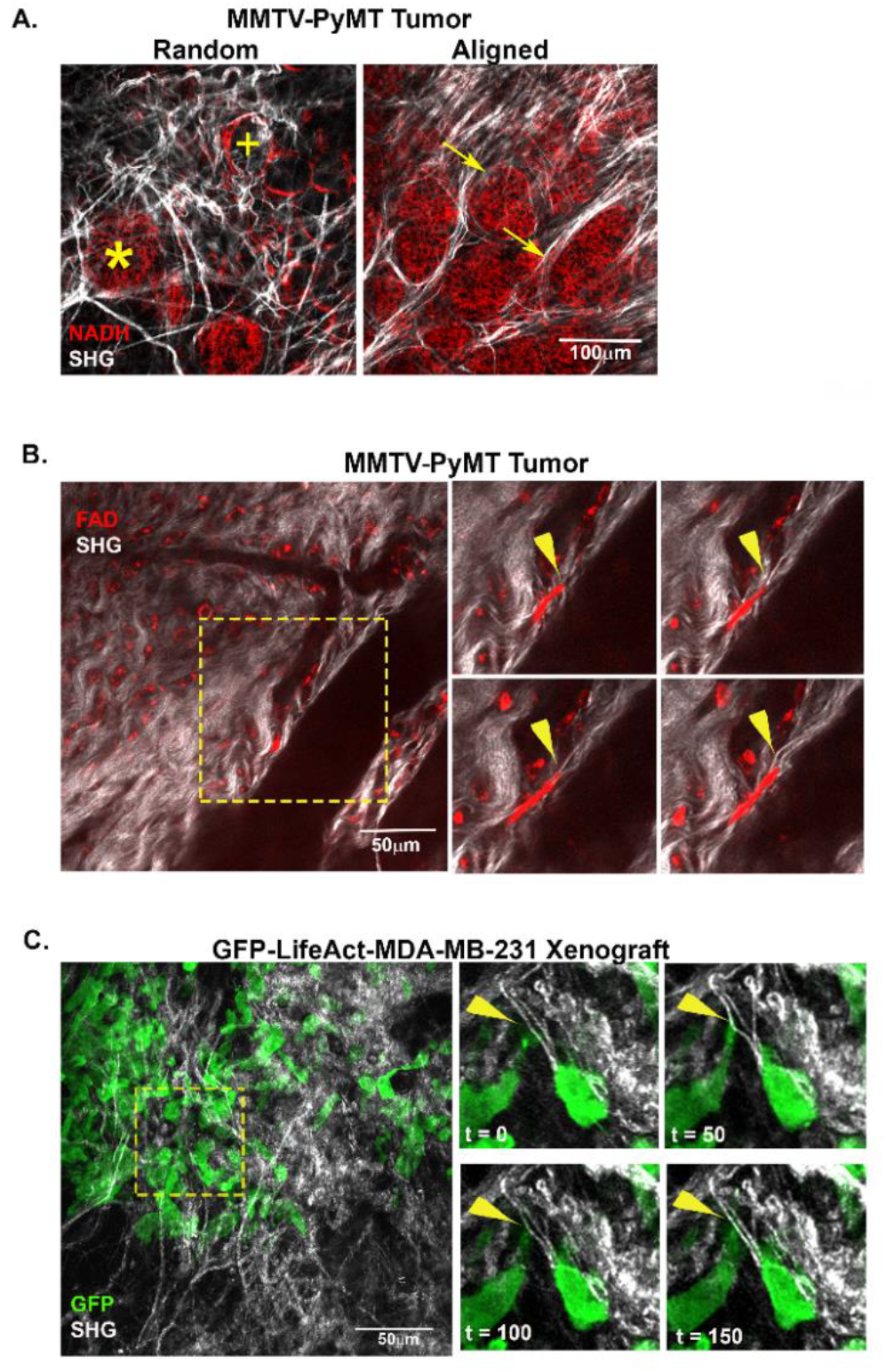
Intravital imaging of cells interacting with and deforming stromal fibers. **(A)** Intravital images of a MMTV-PyMT mouse mammary tumor show both random and align collagen (white) fibers within the tumor stroma. Adipose (+) and tumor (*) cells are visualized within close proximity of each other in regions of random fiber configuration while bundles of aligned fibers can be visualized extending perpendicular to tumor mass (arrow). **(B)** Timelapse movies show cells migrating within tumor microenvironment (TME). The inset (yellow box) highlights a migrating cell deforming an individual fiber (arrowhead). **(C)** Xenograft of GFP-labelled MDA-MB-231 migrating within the TME. The inset (yellow box) highlights a cell extending a protrusion on a collagen fiber and deforming the fiber. Arrowhead indicates point of contact and deformation.

The organization of the ECM in breast cancer is prognostic of poor patient outcome (*7*). Therefore, we examined the behavior of the ECM and its organization during *in vivo* breast cancer cell migration. Once again, SHG was used to monitor the displacement of collagen fibers while endogenous cellular FAD fluorescence (FAD) was used to visualize the actions of resident cells. During the course of these intravital timelapse collections, a vast majority of the cells in the PyMT tumor remained stationary, but a small population of cells could be seen undergoing both amoeboid and mesenchymal migration. In general, migratory cells, but not the stationary ones, deformed individual fibers as they contracted and moved (arrowheads, Figure 1B). Deformations were detected locally at the site of cell attachment and propagated away from the migrating cell within a locally interconnected network of fibers. To better visualize these interactions and achieve greater levels of cell migration under the mammary imaging window, xenografts of MDA-MB-231 stably expressing GFP-LifeAct were also implanted in the mammary gland in parallel experiments (Figure 1C). The behavior of the MDA-MB-231 mirrored that of PyMT tumor cells visualized by endogenous fluorescence with the advantage of being better able to visualize cellular protrusions. Consistent with cells visualized by endogenous fluorescence, these genetically-labeled cancer cells were also visualized displacing collagen fibers within the TME (Figure 1C), thereby confirming that individual cancer cells have the ability to displace and deform individual collagen fibers *in vivo.* These results suggested that the local fiber structure should be taken into consideration when assessing stiffness because individual fibers are not rigid relative to the forces that the cells apply. In fact, the relative rigidity of the fiber is likely dependent on the location and orientation of the fiber relative to the local forces applied to it, evolving dynamically in direction, magnitude and location as the cell migrates. The question remains if we can better characterize the cellular response (i.e. protrusive activity, focal adhesions, etc.) to specific matrix features and how, in turn, do different matrix structures or organizations respond (i.e. deform) to those locally applied forces?

### Cell migration and protrusion dynamics are responsive to the local alignment of collagen fibers

Intravital investigation of the PyMT and the xenograft tumor models clearly demonstrate that migratory cells interact with and deform collagen fibers within the native tumor microenvironment. However, breathing artifacts coupled with lack of control and reproducibility of highly aligned fiber configurations limited in-depth investigation of cell/matrix reciprocity within this system. To better investigate the dynamic interplay between cell behavior and the underlying mechanics governing dynamic fiber deformations in random and aligned matrices, we utilized a mechanical strain device to locally align regions of collagen fibers within a random 3D collagen gel (Figure 2A, Supplemental figure 1A). These aligned regions were the product of permanent deformations produced by the initial straining of the gel and were retained upon the removal of the device. Moreover, they could be produced on-demand, and could easily be distinguished morphologically by their straightened fibers and a high coefficient of alignment as measured by an automated curvelet-based fiber analysis algorithm (*14*). The coefficient of alignment (a measure of similarity between fiber orientations in regions aligned by the mechanical strain device, with a coefficient of 1 representing complete alignment) had a value of 0.75 compared to 0.40 in the unstrained and random controls (Figure 2B). Using this system, we began to systematically investigate how local organization of the collagen fibers defines the cellular behavior and how the underlying mechanics of the matrix exerts a uniquely local influence on the embedded cells as they migrate.

**Figure 2:**
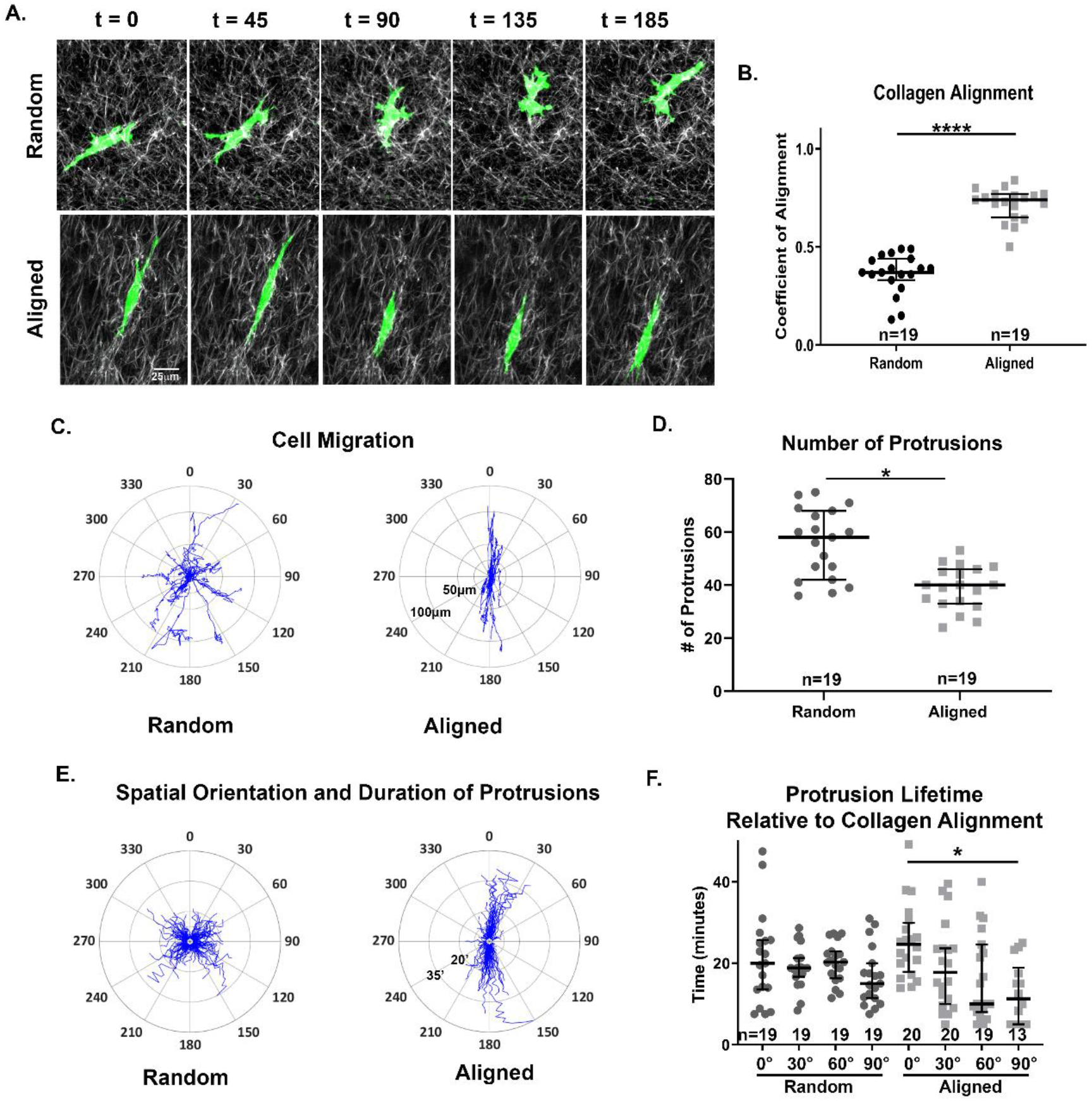
Migratory and protrusive behavior is dependent on fiber organization. **(A)** Timelapse imaging demonstrates morphology and behavior of embedded MDA-MB-231 cells in random and aligned collagen organizations. **(B)** Collagen fiber organization in random and aligned regions of the gel was quantified using the curvelet algorithm, ctFIRE. Windrose plots described the migratory **(C)** and protrusive **(E)** behavior of individual cells in random and aligned matrices. **(D)** Quantification of total protrusions in aligned vs random fiber organizations. **(F)** Protrusion lifetimes were determined and quantified by relating the orientation of each protrustion with respect to the axis of alignment. Medians and interquartile ranges were reported and significance was determined by Wilcoxon rank sum.

Consistent with previous studies, collagen alignment resulted in greater directional migration and persistence (Figure 2C) of cells in 3D gels (*8, 15*). In our aligned gels, cells generally limited their migration path to +/− 30° from the axis of fiber alignment with movements largely restricted to forward and backward motions. In regions of random collagen fiber organization, cell migration paths remained more evenly distributed throughout the matrix and were characterized by random cellular movements with frequent turning motions. (Figure 2C).

We next sought to characterize the effect of collagen alignment on cellular protrusions. We first measured the overall number and spatial distribution of protrusions in both aligned and random matrices using an unbiased, automated approach to quantify cell protrusions (Supplemental Figure 2) (*8*). In cells localized to regions of aligned fibers, the median number of protrusions significantly decreased compared to cells in random control matrices (59 to 40, respectively). More in-depth spatial and temporal analysis of individual protrusions revealed a clear pattern between those protrusions extended along the axis of alignment and those oriented perpendicular to it. While protrusions in random collagen matrices were uniformly distributed with respect to their frequency, spatial distribution, and duration, in aligned matrices, the number of protrusions were once again limited to the axis of fiber alignment (Figure 2e). Individual protrusions extended within 15 degrees of the axis of alignment were both more frequent and significantly longer-lived (24.7 min median lifetime) than those extended perpendicular to it (11.3 min median lifetime) (Figure 2F). It is worth noting that although reduced in number and longevity, protrusions perpendicular to the cell and fiber axis could still be visualized interacting with the matrix but were quickly retracted with contractions not being sustained.

### Focal adhesion orientation mirrors collagen fiber organization

Focal adhesions (FA) likely regulate aspects of the protrusive behavior observed in 3D matrices. FA exert cellular forces on the matrix and respond to the amount resistance, thereby providing the cell with intrinsic and immediate feedback of local extracellular cues. In 2D, FA size correlates with increased mechanical force (*16*), and larger, elongated focal adhesions are often associated with increased traction forces along the long axis of the FA (*17*). For the purpose of this study, we used immunofluorescence to examine the distribution of phosphorylated FAK and vinculin as surrogates for mature, force-generating FA with respect to collagen fibers and matrix organization. FA positive for both pFAK and vinculin were located along collagen fibers in both random and aligned collagen matrices (Figure 3A). A majority of these FA were distributed in the distal cell protrusions, but some FA were still found at the periphery of the cell body, where cell-fiber contact occurred where the cell was in contact with a collagen fiber (arrows). Surprisingly, when we examined the overall size of pFAK and vinculin plaques, the average FA size was the same in both types of matrices (Figure 3B, 3C). However, when we specifically determined the aspect ratio (Figure 3D, 3E) of pFAK and vinculin positive FA with respect to the aspect of alignment, the median aspect ratio of FA oriented parallel to the axis of alignment was increased (i.e., elongated) as compared to FA that formed perpendicular to the axis of alignment (1.50 vs 1.38 for pFAK, 1.56 vs 1.45 for vinculin, Figure 3D, 3E). In random matrices, there was no difference in the aspect ratio between similar orthogonal directions. Moreover, in cases where the cell body and aligned fibers shared a common axis, prominent plaques of vinculin could be identified associated with increased SHG signal due to the accumulation of collagen (arrowhead, Figure 3A). This increase in SHG signal is consistent with enhanced cell generated forces or contractility due to increased directional resistance or stiffness from aligned fibers. This accumulation was generally not observed in random fiber configurations.

**Figure 3:**
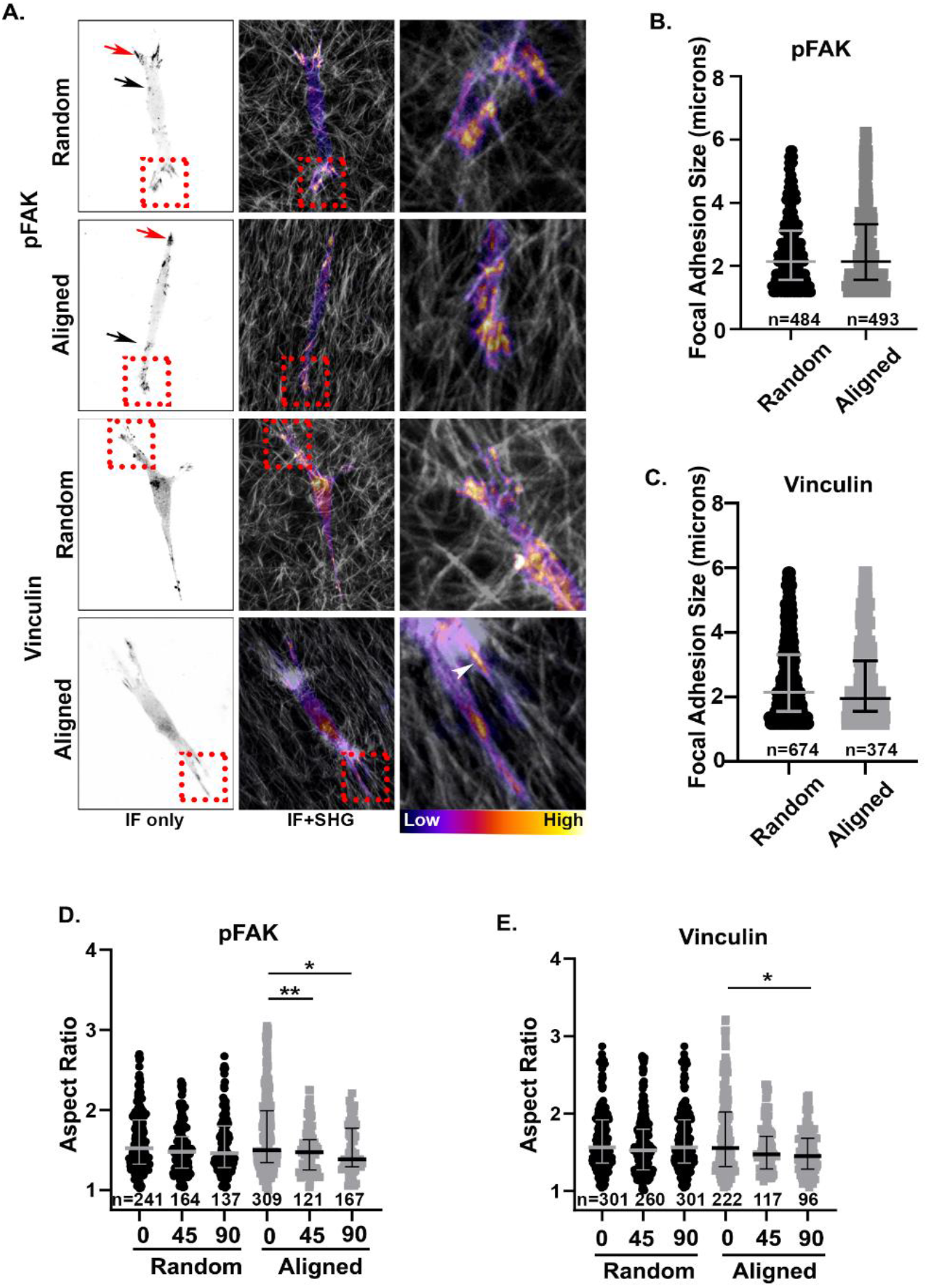
Focal adhesions (FA) elongate along the axis of alignment. **(A)** Immunofluorescence of pFAK and Vinculin identify the distribution of focal adhesions in MDA-MB-231 cells embedded in aligned and random collagen matrices (white). FA plaques contact individual collagen fibers (black arrows) primarily localized to distal protrusions (red arrows), but are also found witinin the cell body. Magnified regions (red box inset) highlight FAs localized within the cell protrusion. Higher magnification shows an intensity image of vinculin and pFAK expression organized along individual collagen fibers. **(B,C)** Quantification of vinculin and pFAK size/cell in random vs, aligned conditions. Quantification of the aspect ratio of pFAK **(D)** and vinculin **(E)** relative to the axis of fiber alignment. Medians and interquartile ranges were reported and significance was determined by Wilcoxon rank sum.

### Orientation of fibers relative to applied forces affects local stiffness

Fibrous networks like those of the ECM stiffen under increasing strain (*8, 18, 19*), and are stiffer along the axis of alignment (*8*). This understanding, however, is derived from macroscopic experiments, and it remains unclear how matrix stiffness translates to the microscopic scale where cells interact with individual or small heterogenous networks of fibers. At the cellular scale, local mechanical properties are likely better described by the collective action of relatively small collections of fibers. Collagen and other ECM fibers have small diameters relative to their length resulting in a small bending stiffness and negligible ability to resist orthogonally applied forces (*20, 21*). Due to this property, cellular perception of stiffness among equivalent fibers would also depend on the orientation of the fiber relative to the vectors of forces exerted. In other words, a FA exerting a force perpendicular to the fiber would immediately deform the fiber with little resistance. However, a focal adhesion applying force parallel to the fiber would experience far greater resistance to deformation (i.e. greater stiffness) resulting in more sustained tension across the plasma membrane to the cellular contractile machinery.

To quantify the difference in stiffness between orthogonal directions in random and aligned matrices, we used an acellular approach utilizing microspheres made of temperature-sensitive poly N-isopropylacrylamide (pNIPAAm). These cell-sized microspheres were embedded within 3D collagen gels and covalently attached to the fibers. Upon a precise increase in temperature, the microspheres apply controlled cell-scale contractions to the matrix. This approach has been successfully used to investigate ECM micromechanics (*22–24*), and its advantages are that it can produce cell-scale deformations in 3D collagen matrices without modifying the matrix (e.g. by depositing additional matrix or degrading the existing one). Additionally, advances in this methodology also allow for the pNIPAAm microspheres to be calibrated to moduli of known stiffness thereby providing a quantitative measure of the local modulus at different locations within the 3D matrix (*24*). By using these techniques, it is possible to quantify cell-scale differences in local matrix stiffness due to changes in fiber architecture through measurements of the resulting matrix deformations and changes in the aspect ratio of the particles.

Calibrated pNIPAAm microspheres were embedded into three different types of 3D collagen gels: the unstrained random controls, aligned gels with the strain device left in place, or aligned with the strain device removed to ensure the device was not restraining the matrix or transferring forces. At room temperature, the microspheres in all conditions were rounded and remained unstrained (Figure 4a). Upon elevating the temperature, the microspheres contracted isotropically, thereby exerting equivalent radial forces on the covalently attached collagen fibers. In the random collagen gels, the microspheres contracted uniformly; maintaining a spherical shape, indicating that the matrix stiffness was symmetrical in all directions. The microspheres also contracted in both aligned matrices. However, the microspheres were noticeably elongated along the axis of fiber alignment, indicating an effect of direction on the stiffness within the matrix. Notably, there was no significant difference between the aspect ratios of aligned matrices with or without the strain device, only between the aligned and random controls.

**Figure 4:**
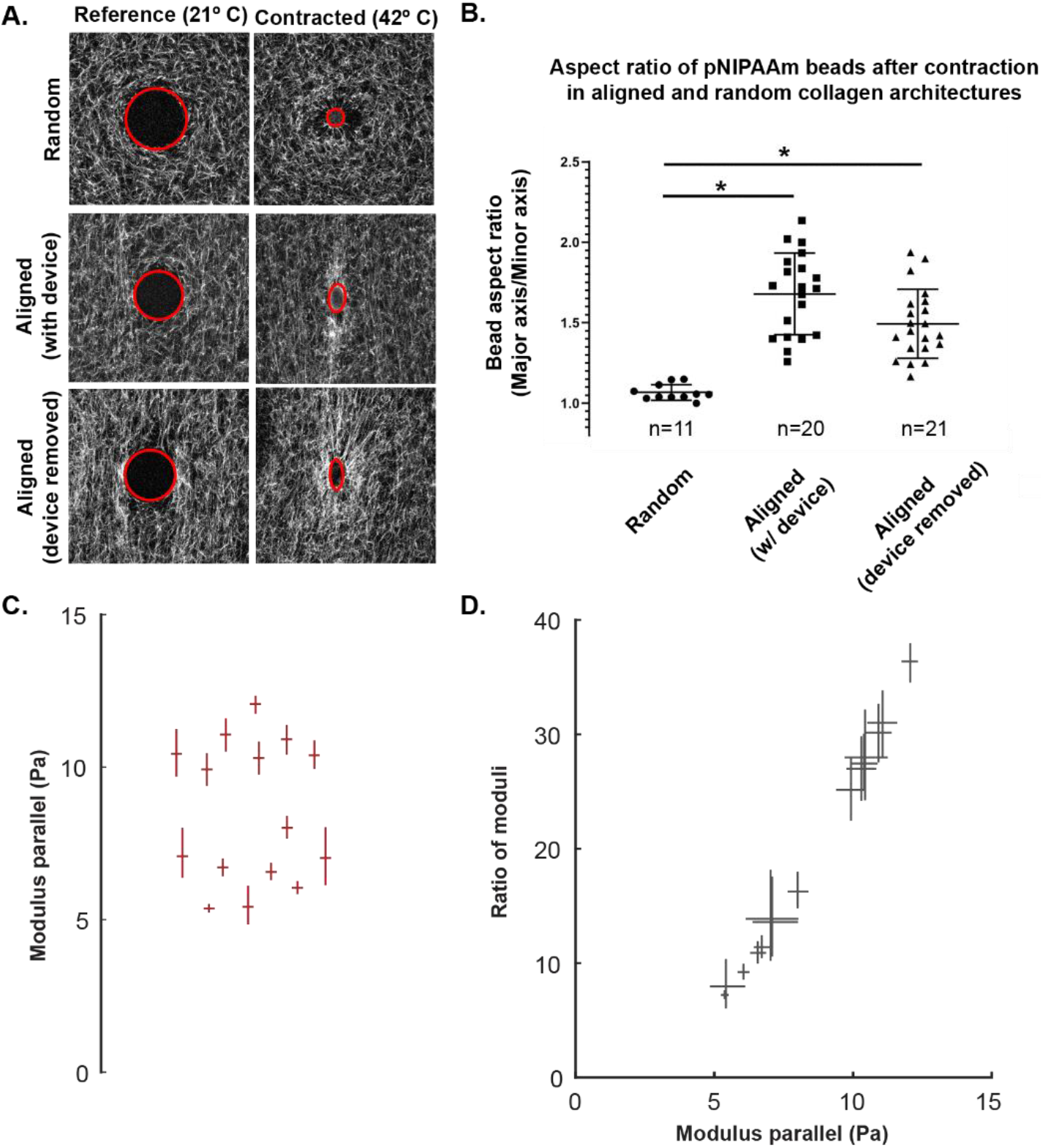
pNIPAAm beads measure local modulus in orthogonal directions in random and aligned matrices. **(A)** Prestrained reference and post-contraction images of fluorescent collagen with pNIPAAm beads (edge of bead defined by red line) show anisotropic moduli across orthogonal directions with the 3D matrix. **(B)** Quantification of the bead aspect ratio. Means are reported with standard deviation, and significance was determined by one-way ANOVA. **(C)** Calculation of the local modulus parallel to axis of alignment. **(D)** Plot of ratio of the parallel/perpendicular moduli vs the parallel modulus. For panel C, each horizontal line equals calculated value while vertical bars are 95% CI; for panel D, both lines indicate the 95% CIs and the intersection equals the calculated value

To quantify the relative magnitudes of stiffness between parallel and perpendicular directions relative to fiber alignment, we used the measured strains of the calibrated pNIPAAm microspheres to calculate local modulus. By adapting a previous technique (*24*) we measured the modulus in orthogonal directions of an aligned matrix, thereby providing a directional stiffness and a relative microscale measure of how much stiffer the gel is parallel than perpendicular to fiber alignment (for details, see Supplemental Note). The mean cell-scale modulus along the axis of alignment was calculated to be 8.5 Pa, which is approximately 16-fold greater the modulus in directions perpendicular to fiber alignment (Figure 4c). In addition, we compared the relative measures of stiffness through the ratio of their strain and therefore moduli across directions. From this analysis, the modulus in the direction of alignment could be as high as 25 to 30 times the stiffness of the perpendicular axis (Figure 4d). When this result is combined with the measurements of aspect ratios from the pNIPAAm beads, it reveals the effects of local structures are significantly greater than one would expect. To demonstrate this point, examine both the aspect ratios of pNIPAAm beads (parallel:perpendicular) and the ratio of moduli (parallel:perpendicular) from regions of aligned collagen fibers. The median aspect ratio of the contracted beads in aligned regions was 1.71 (Figure 4b), but the difference in stiffness across axes is not a factor of 1.7; it is a factor of 25 (Figure 4d). This clearly demonstrates that local organizations and alignment provide a surprisingly sizeable and often underestimated directional cue.

### Collagen fiber alignment biases matrix deformations and the propagation of fiber displacements

In addition to the differential stiffness cues that result from the orientation and organization of collagen fibers, matrix deformation in random and aligned conditions may also be different. This could affect reorganization of the ECM and the subsequent cellular mechanosensing. To investigate this further, we used digital image correlation (DIC) (*25*), a method that provides accurate, full-field measurements of displacements using only the fibers themselves, and embedded contractile pNIPAAm microspheres to induced localized deformations within the collagen gel (Figure 5a). In the random matrices, the displacement pattern was generally uniform and circular (Figure 5a). In the aligned gels, however, the displacement fields were noticeably warped and elongated along the axis of fiber alignment (Figure 5a). That is, the displacements propagated further along the axis of alignment than perpendicular to it. This was counterintuitive, as the spherical pNIPAAm beads contracted less on this axis parallel to alignment and become elongated due to the anisotropic moduli of the surrounding matrix. To quantify this observation, we drew 60 equally spaced lines radiating around the contractile bead (red, blue and gray traces, figure 5b) and extending toward the edge of the image. The DIC displacements along each line were extracted and plotted over distance. In the aligned matrices, the traces along the axis of alignment (figure 5b, shown in red trace) show a slow decrease in matrix displacements over distance. Conversely, displacements perpendicular to the axis of alignment (Figure 5B, blue) decrease rapidly with steeper slope, and clearly differed from the traces parallel to fiber alignment. In random controls, orthogonal measurements demonstrate that radial displacements were approximately equivalent over distance, indicating no difference in mechanical behavior.

**Figure 5:**
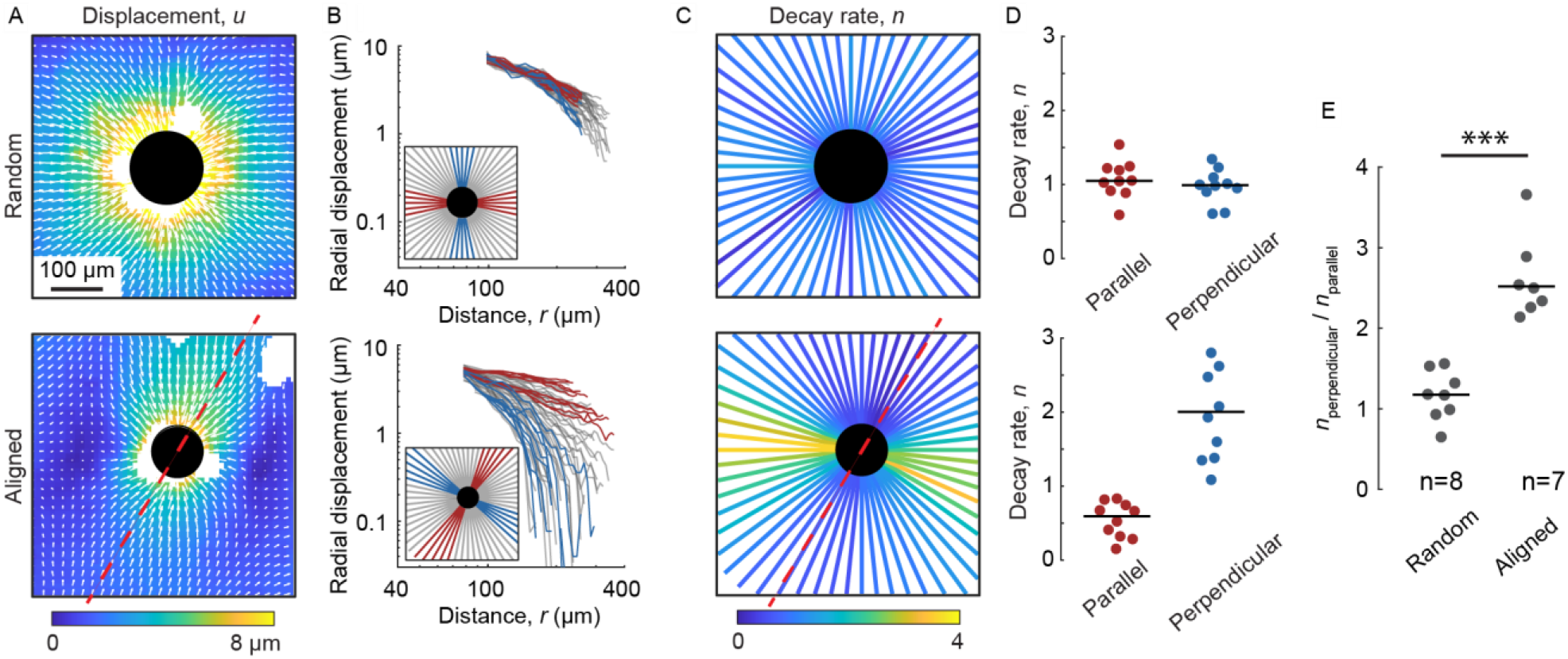
The alignment of collagen fibers imparts directionality on the propagation on contraction-induced displacements. **(A)** Displacement maps from random and aligned labelled collagen gels embedded with pNIPAAm beads. The axis of alignment is represented by the red dashed line. **(B)** Radial traces were extracted from the displacement plots and plotted against the distance from the contractile particle. A key describing the location of the individual trace is located in the lower left corner of each plot. Red lines represent traces extracted along the axis of alignment while blue lines represent traces perpendicular to alignment. Gray lines represent non-orthogonal angles. **(C)** Map and **(D)** quantification of calculated decay rates, *n*, for extracted traces. **(E)** Quantification of the ratio n_perpendicular_/n_parallel_ in multiple beads in both random and aligned matrices. Lines indicate medians; significance was determined with a Wilcoxon rank sum test.

To further quantify and characterize the fiber displacement properties observed in the traces from aligned and random matrices (figure 5b), we fit the radial displacements *u* to distance from the center of the pNIPAAm microsphere *r* to the equation *U*_r_ = A*r*^−n^ as in previous studies (*10, 23*). The fitting variable *n* represents how quickly the displacements diminish over distance. For reference, displacements in linear elastic materials like polyacrylamide decay very rapidly with *n* = 2 (*26*). Fibrous, biological materials like collagen gels decay much more slowly with *n* ≈ 1. Consistent with previous work and theoretical models (*10, 23, 27*), the decay rates in matrices with random fiber organization decayed slowly with values of *n* near to 1 (Figure 5 c,d). Comparisons of *n* in orthogonal directions showed no significant difference in decay rates (figure 5d), and the ratio of *n* in two orthogonal directions equaled 0.94, indicating the matrices were essentially isotropic. In contrast, in aligned matrices the decay rate *n* was no longer uniform through the material. Perpendicular to the axis of alignment, the decay rate increased to values greater than 2 (i.e. faster decay of displacements than in linearly elastic materials), while parallel to the axis of alignment the decay rate decreased to 0.5 (i.e. slower than in random collagen networks) (Figure 5c, d). Under these conditions the ratio of the decay rate in directions perpendicular and parallel to fiber alignment (*n*_perpendicular_/*n*_parallel_) increased to 3.38, representing a greater than 3-fold increase in the decay rate (figure 5d). This process was repeated for multiple microspheres in both random and aligned matrices. In random matrices, the ratio of *n* in orthogonal directions was centered around one; however, for microspheres embedded in aligned gels the ratio was greater than 2, indicating a significant difference in decay rate due to matrix organization (Figure 5e). Similar to the findings from the pNIPAAm bead aspect ratios and ratios of modulus in aligned collagen gels, the effect of alignment on the decay rates is significantly greater than expected, and potentially represents sizeable cue to the cell. Moreover, this finding suggests that propagation of displacements in fibrous materials like the ECM is not homogeneous but rather variable and dependent on the organization of the fibers.

To validate that similar deformation and decay patterns arise due to cellular behavior and individual protrusions within random and aligned collagen architectures, MDA-MB-231 cells were embedded in collagen gels, and the displacements that they produced were measured with DIC. From this analysis, discrete pulses of displacements could be visualized propagating from individual protrusions instep with cell-induced contractions. Due to the pulsatile nature of these protrusions, an equivalent technique and sampling to that of the pNIPAAm beads was done on the average of all cell induced displacements over time. We observed that cell-induced displacements in random fiber organizations exhibited no discernable differences in the decay rates between orthogonal directions (figure 6 A and B). In aligned matrices, however, the decay rates along the axis of alignment (red trace, figure 6) once again differed from the decay rates of those perpendicular to the axis of alignment (blue trace, figure 6) mirroring the results of the pNIPAAm microspheres. To better visualize this behavior and account for variability of cell shape, we plotted the displacements of the matrix relative to the cell perimeter by circumferentially extracting the displacements from the edge of each cell and incrementally expanding outwards to generate an average matrix displacement map with respect to the cell edge. This data was then plotted on polar coordinates to give a 360-degree distribution of displacements over time. From this analysis, we observe many similar patterns with the pNIPAAm beads. That is, in random matrices, the displacements were more uniform throughout while in aligned matrices, there was a clear difference in behavior between displacements generated on axis and those generated perpendicular to it. The displacements located parallel to the axis of alignment were smaller yet propagated over a longer distance, while those generated perpendicular to the axis of alignment exhibited greater magnitude but a shorter range. This supported our findings from both the pNIPAAm bead experiments that the alignment of collagen fibers not only affects the directional stiffness, but also how far those displacements propagate.

**Figure 6:**
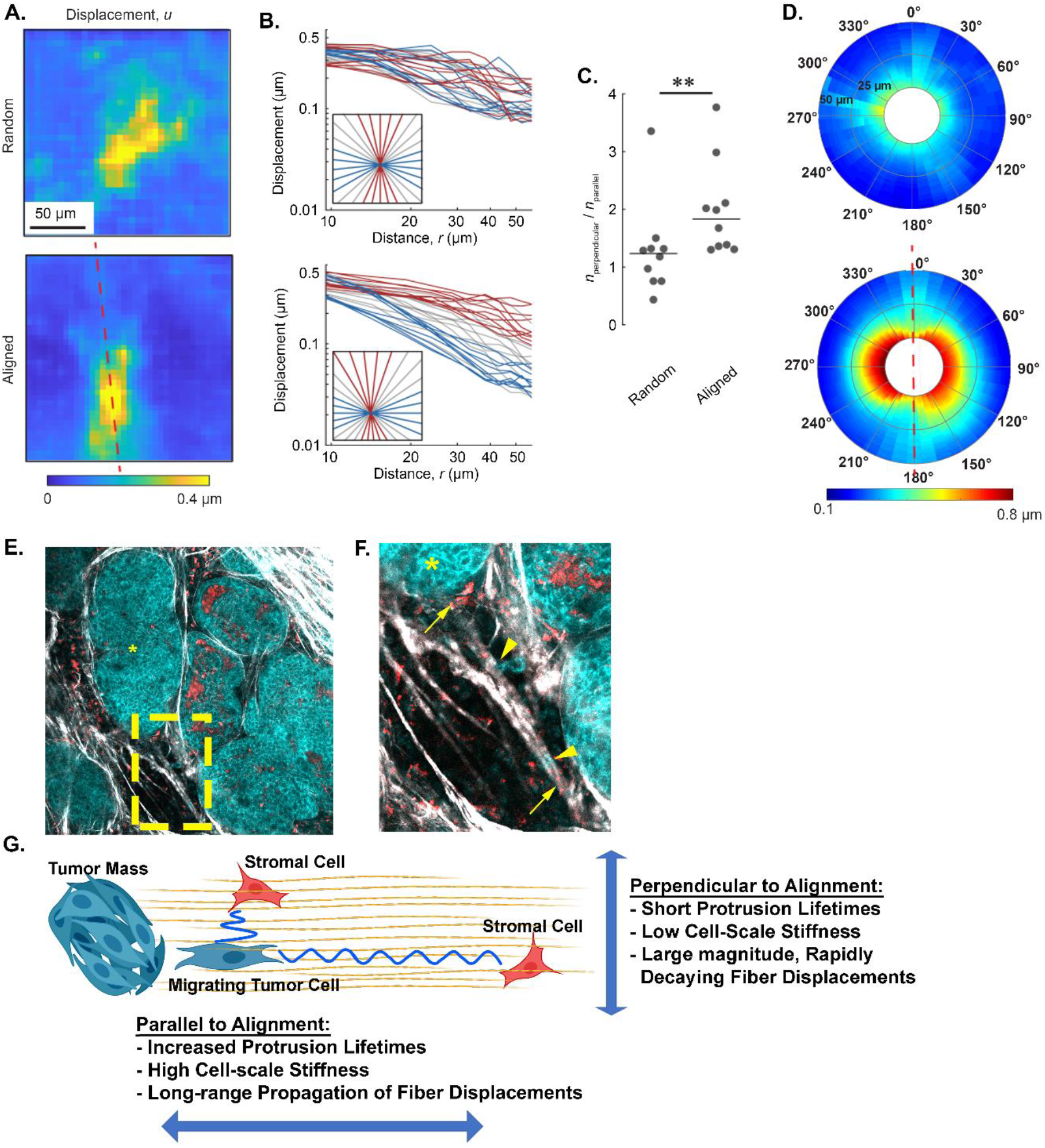
Fiber organization affects the propagation of cell-induced displacements in fibrous ECM environments and impacts cell mechanosensing. (A) Displacement maps from random and aligned labelled collagen gels embedded with MDA-MB-231 GFP-LifeAct. The axis of alignment is represented by the red dashed line. (B) Radial traces were extracted from the displacement plots and plotted against the distance from the contractile cell. A key describing the location of the individual trace is located in the lower left corner of the plot. Red lines represent traces extracted along the axis of alignment while blue lines represent those that extend perpendicular. Gray lines represent non-orthogonal angles. (C) Quantification of the ratio n_perpendicular_/n_parallel_ from multiple cells in both random and aligned matrices. (D) Displacements were circumferentially extract around the perimeter of a cell and plotted on 360° polar plots to describe the pattern of average displacements relative to the cell perimeter. The axis of alignment was represented by the red dashed line (A, D). (E) A representative field of view taken from an intravital mouse mammary tumor shows multiple cell types interacting within random and align collagen (white) fibers in proximity to tumor masses (cyan, *, E). The tumors and disseminating tumor cells are visualized with NADH endogenous fluorescence (cyan) while stromal cells could be visualized with with FAD autofluorescence (red). Higher magnification (yellow inset, F) of the tumor highlights stromal cells and stromal cells potentially interacting with each other and local fibers. (G) A schematic of how fiber organization may facilitate or bias mechanical cues used in mechanosensing. Displacements generated along the axis of alignment propagate further than those that might propagate perpendicular to alignment. This might allow cells to disproportionately sense other cell types at greater distance along the axis of alignment. In panel C, lines indicate medians; significance was determined with a Wilcoxon rank sum.

## Discussion

This study aimed to capture a more comprehensive perspective of the cell-matrix mechanosensing system, including both the physical cues that direct and orient cellular adhesions to efficiently organize and apply forces, and the characteristics of the subsequent cell-scale matrix deformations that may further instruct cellular behavior. To accomplish this, we used intravital imaging to evaluate the level of reciprocity between the cells and the matrix in vivo. Then, we developed an *in vitro* model where we could readily investigate the mechanics of the underlying matrix. From the intravital studies, we were able to visualize the heterogeneity present within a physiological ECM where both random and aligned fiber configurations were abundant. Moreover, from these images, individual cells could be seen displacing individual fibers as they applied contractile forces necessary for locomotion. These findings suggested that we consider the resultant deformation from applied forces when we assess stiffness in physiological tissues. We then took a closer look at commonly observed fiber organization within a 3D *in vitro* collagen gel. In an aligned fiber configuration, cells migrated more persistently with fewer, longer-lived protrusions localized to the axis of fiber alignment. Similarly, FA plaques could be visualized co-localizing with collagen fibers. The FA organized/oriented along the axis of alignment had greater aspect ratios compared to their perpendicular counterparts, suggestive of local anisotropy in stiffness. To further investigate the local stiffness with respect to fiber alignment, we used calibrated pNIPAAm beads and determined that the local modulus, while variable, could be up to 30 times stiffer along the axis of alignment than perpendicular to it. We also observed that in addition to increasing the local anisoptropy in modulus of the matrix, fiber alignment supports the long-range propagation of displacements while diminishing those same signals that propagate perpendicular to that fiber organization. This has the implication that mechanosensing potential is not uniform throughout the ECM, but rather the organization of the matrix may either bias or shield physical signals, thereby restricting or constraining biophysical communication within the microenvironment.

One of the more important understandings to come out of this study is how local fiber organization might modulate the relationships between the mechanical cues and cell behavior in the 3D environment. Various mechanical cues comprise the heterogeneity found in the extracellular matrix, including contact guidance, durotaxis, and physical confinement. A challenge that remains is how to unravel and define these integrated cues to better understand the cellular response. Intuitively, it makes sense that interdependencies between mechanical cues exist. After all, the growth of focal adhesions is restricted to individual fibers which all have their own unique mechanical properties that vary depending on the orientation of the forces that are applied. While that concept is generally appreciated, the magnitude to which local architecture of the fibrous matrix impact these mechanical cues is hard to decouple and characterize. Physiological, fibrous networks like the ECM have numerous non-linear properties that may provide a platform to rapidly amplify or diminish mechanical cues. Our study highlights non-linearity in the matrix by identifying a dramatic difference in cell-scale modulus driven by collagen fiber organization, which is associated with an increase in protrusion longevity, FA elongation and directed cell migration. Thus, the properties of the matrix may play a large role in the many biphasic or nonlinear behaviors exhibited by cells.

An added dimension of cell-matrix interactions highlighted in our study is how the local matrix deforms in a 3D microenvironment due to applied cellular forces during migration and invasion. Recently, macrophages were shown to be attracted to the tugging action of myofibroblasts at the wound site to facilitate healing (*12*). It was proposed that sensing mechanical actions by the macrophages would have the advantage of sensing the active myofibroblasts at greater distances than those achievable by chemical gradients. While that study focused on wound healing and the interactions of macrophages and myofibroblasts, it is certainly plausible that other fibrotic conditions or cell types could use similar behavior to sense specific dynamic cues or structures at significant distances. During cancer progression, collagen fibers aligned perpendicular to the tumor stromal boundary (TACS-3) are predictive of poor patient outcome (*7*). It is also known that aligned fibers increase persistent cancer cell migration and are enriched in macrophages (*8, 28, 29*). The mechanisms of cells trafficking to regions of aligned collagen fiber in the tumor microenvironment are not yet clear, but understanding how cell forces propagate through the matrix, specifically within aligned collagen fibers, might shed light on an additional biophysical mechanism of cell-cell communication. For example, tumor cells have been shown to pair with macrophages to facilitate tumor cell intravasation into blood vessels. In these situations, reciprocal chemokine signaling facilitates macrophage-tumor cell trafficking along collagen fibers (*30, 31*), but the organization of matrix may play a significant role in initiating the pairing of these cells by signaling through long-range fiber displacements. This process may be further leveraged or amplified because regions of local alignment may bias the detection of the tugging cell by the homing macrophage at a much greater distance than those that might be perpendicular to this alignment (Figure 6G). Conversely, any cell positioned perpendicular to the axis of alignment would be effectively shielded to those same displacements.

In summary, a key principle of this study is that mechanical cues in physiological 3D environments are heterogeneous and highly dependent on the local organization of the fibers. The organizational and mechanical details of local microenvironment are truly paramount. Our work has identified a significant difference in several cell-scale mechanical cues (stiffness, the decay rate of collagen fiber displacements, etc.) with respect to the organization of collagen fibers. We are just beginning to scratch the surface of defining how mechanical properties of the matrix impact the reciprocity in cell behavior and cell communication within the microenvironment. Future studies are needed to investigate how variable compositions or specific features of the matrix will further enhance or diminish these cues, and how dynamic signaling plays a role in these processes.

## Materials and Methods

### Cell culture

MDA-MB-231 (ATCC) cell lines used in this study were maintained in DMEM (1g/L glucose, L-glutamine, 110mg/l Sodium Pyruvate) with 10% FBS under 5% CO_2_. The LifeAct-GFP construct (kind gift from Dr. Maddy Parsons, King’s College London) was used to create stable cell line as previously described (*8*).

### In vivo imaging experiment

B6 mice expressing the MMTV-PyMT transgene were raised for 8-10 weeks until a palpable mass was detected. SCID mice (Jackson Lab) were injected with 2.5×10^6^ MDA-MB-231 cells expressing LifeAct-GFP over R4 mammary fat pad. Cells were allowed to grow until palpable (10-14 days). In both the PyMT and MDA-MB-231 xenografts, once the tumor was palpable and approximately 5mm in diameter, a Mammary Imaging Window (MIW) was surgically implanted over the tumor mass (*13*). Tumor was then allowed to grow into the window for another 3-5 days until imaging.

### Imaging and microscopy

All imaging was done using a Bruker Ultima Multiphoton Microscope equipped with a Coherent Chameleon Ti-Sapphire laser and Hamamatsu R3788 multi-alkali photomultipliers. For live cell protrusion dynamics, laser imaging was conducted at 850nm with a Zeiss 20x NA 1.0 objective lens, and GFP(525/70) and RFP(565LP) filter cubes. For focal adhesion analysis, imaging was conducted at 890 nm using SHG(450/40) and GFP(520/40) filter cube and Nikon 40x 1.15 NA LWD objective lens. For measuring anisotropic modulus and field displacements of Ax-488 collagen gels with pNIPAAm microspheres, the laser was tuned to 890nm and collected with a Nikon 20x NA .45 objective and the GFP filter cube.

### Collagen polymerization

Rat-tail collagen (Corning Inc.) was labelled with Alexa-594 or Alexa-488, neutralized, and polymerized as previously published (*32*). Briefly, cell seeded collagen gels were made by mixing fluorescent labeled collagen with unlabeled collagen at 1:10 ratio. The solution was neutralized using HEPES buffer and then 2.5×10^6^ cells in media were added to the solution for a final collagen concentration of 2mg/ml. The collagen/cell solution was then placed on a thermoelectric plate (CP-061HT, TE technology) with a TC-720 temperature controller (TE technology) set at 21° C and allowed to polymerize for 60 minutes before placing in a 37° C incubator for 15 min. Gels were then strained and allowed to incubate overnight before imaging.

### Mechanical alignment of collagen gels

The mixture of neutralized collagen with either beads or cells were cast in the bottom of 20mm glass bottom dish (CellVis). After polymerization, the gel was mechanically aligned by a strain device adapted from Vader et al (*33*). Briefly, lids from 35mm dishes were modified to have two cantilevers with cushioned ends that apply point loads to the gel producing localized strains between the two cantilevers. The cantilevers were 2-3 mm apart and strained another 2-3 mm gels were strained until alignment of the fibers could be observed by a dissecting microscope with a 10X objective.

### Quantification of protrusion dynamics

Protrusions were identified using Matlab script as described in Riching et al (*8*). In order to quantify protrusion dynamics, protrusion dots as identified were calculated for each time frame in cell migration movie which then was transferred to dots in a blank tiff file (Supplemental figure 2). A Gaussian filter was then applied using FIJI for each image. Then a particle tracking plugin from MosaicSuite in FIJI was utilized to track each dot. Dots that were tracked were considered one protrusion.

### Immunofluorescence for focal adhesion identification

Cells were fixed with 4% PFA in PBS for 45 min, permeabilized using 0.2% Triton-X in PBS, and subsequently blocked overnight using 1% BSA in PBS. Cells were then treated with primary antibody at 1:200. Anti-Phosphorylated FAK 397 (Invitrogen, catalogue #44-624G:) or Anti-vinculin (Sigma, cat #v4505:). After extensive washing, cells were incubated in secondary Alexa-488 anti-rabbit or anti-mouse antibody (Thermo Fisher Scientific) at 1:500 dilution.

### Quantification of focal adhesions

Images were gathered using z-stacks at 1 um step size using a Nikon 40x NA 1.15 LWD objective lens. For FA identification, cell images from fluorescent channel were z projected using maximum and average intensity using FIJI. Average z projection was then subtracted from maximum z projection. The subsequent image had a background subtraction with a rolling ball radius of 5 pixels applied. Images were then thresholded so that only the top 0.5% pixels were included using FIJI threshold and a binary mask was applied. Using Analyze Particles function in FIJI, each FA coordinate was recorded and then transferred to MATLAB. In MATLAB, centroid of FA and area were calculated using polygon function. Length and width of FAs were calculated by using fit ellipse function in MATLAB for each FA (supplemental figure 3).

### Measurement of anisotropic modulus and full-field displacements with pNIPAAm microspheres

To measure modulus, experiments were performed following the method of Proestaki et al. (*24*) with an additional analysis to consider the effects of anisotropy (see Supplemental Note). First, the thermal strain of the contractile beads in no matrix was measured. Then, the bulk modulus of the contractile beads was calibrated by embedding in polyacrylamide of a known modulus, measuring particle contraction upon a temperature change, and applying Eshelby’s solution (*26*). After calibration, the beads were treated with Sulfo-SanPAH and embedded in collagen gels. Upon inducing local contraction of the beads, the strains in directions parallel and perpendicular to fiber alignment were measured, allowing for computation of the local modulus of the network as described in the Supplemental Note. For measurement of full-field displacements, an image of an unstrained reference state at the polymerization temperature was collected before increasing the temperature to induce localized contraction. Digital Image correlation was performed as previously described(*22, 23*) using the fast-iterative digital image correlation (FIDIC)(*25*).

### Statistical analysis

GraphPad Prism v7.04 was used for generation of graphs and statistical analysis All studies were performed in at least 3 independent experiment containing multiple independent events. Normality of the data was tested using a Shapiro-Wilk test, and for statistical differences between multiple comparisons, a Kruskal-Wallis test was used.

## Acknowledgments

This work was supported by the National Institutes of Health to SMP (RO1 CA142833, RO1 CA216248, and RO1 CA206458) and the National Science Foundation to JN (NSF CMMI-1749400). The authors would like to thank Drs. Anna Huttenlocher and María Virumbrales-Muñoz for helpful discussion and careful review of the manuscript.

## Author contributions

J.S designed, collected, and analyzed intravital and *in vivo* cell experiments.

D.I. collected intravital experiments.

M.P. analyzed local pNIPAAm modulus.

J.N. analyzed fiber displacements and calculated local modulus

B.B designed and collected pNIPAAm experiments, wrote and edited manuscript.

S.P. project administration, edited manuscript, and funding acquisition.

## References and Notes

1. M. J. Paszek, N. Zahir, K. R. Johnson, J. N. Lakins, G. I. Rozenberg, A. Gefen, C. A. Reinhart-King, S. S. Margulies, M. Dembo, D. Boettiger, D. A. Hammer, V. M. Weaver, Tensional homeostasis and the malignant phenotype. Cancer Cell. 8, 241–254 (2005).

2. D. E. Discher, P. Janmey, Y. L. Wang, Tissue cells feel and respond to the stiffness of their substrate. Science (80-.). 310, 1139–1143 (2005).

3. P. P. Provenzano, D. R. Inman, K. W. Eliceiri, P. J. Keely, Matrix density-induced mechanoregulation of breast cell phenotype, signaling and gene expression through a FAK-ERK linkage. Oncogene. 28, 4326–4343 (2009).

4. K. Hu, L. Ji, K. T. Applegate, G. Danuser, C. M. Waterman-Storer, Differential transmission of actin motion within focal adhesions. Science (80-.). (2007), doi:10.1126/science.1135085.

5. M. L. Gardel, I. C. Schneider, Y. Aratyn-Schaus,, C. M. Waterman, Mechanical Integration of Actin and Adhesion Dynamics in Cell Migration. Annu. Rev. Cell Dev. Biol. (2010), doi:10.1146/annurev.cellbio.011209.122036.

6. L. M. Owen, A. S. Adhikari, M. Patel, P. Grimmer, N. Leijnse, M. C. Kim, J. Notbohm, C. Franck, A. R. Dunn, A cytoskeletal clutch mediates cellular force transmission in a soft, three-dimensional extracellular matrix. Mol. Biol. Cell (2017), doi:10.1091/mbc.E17-02-0102.

7. M. W. Conklin, J. C. Eickhoff, K. M. Riching, C. A. Pehlke, K. W. Eliceiri, P. P. Provenzano, A. Friedl, P. J. Keely, Aligned collagen is a prognostic signature for survival in human breast carcinoma. Am. J. Pathol. 178, 1221–1232 (2011).

8. K. M. M. Riching, B. L. Cox, M. R. R. Salick, C. Pehlke, A. S. S. Riching, S. M. Ponik, B. R. R. Bass, W. C. C. Crone, Y. Jiang, A. M. Weaver, K. W. W. Eliceiri, P. J. J. Keely, 3D collagen alignment limits protrusions to enhance breast cancer cell persistence. Biophys. J. 107, 2546–2558 (2015).

9. A. Ray, O. Lee, Z. Win, R. M. Edwards, P. W. Alford, D. H. Kim, P. P. Provenzano, Anisotropic forces from spatially constrained focal adhesions mediate contact guidance directed cell migration. Nat. Commun. 8, 1–17 (2017).

10. J. Notbohm, A. Lesman, P. Rosakis, D. A. Tirrell, G. Ravichandran, Microbuckling of fibrin provides a mechanism for cell mechanosensing. J. R. Soc. Interface. 12 (2015), doi:10.1098/rsif.2015.0320.

11. A. D. Doyle, K. M. Yamada, Mechanosensing via cell-matrix adhesions in 3D microenvironments. Exp. Cell Res. 343, 60–66 (2016).

12. P. Pakshir, M. Alizadehgiashi, B. Wong, N. M. Coelho, X. Chen, Z. Gong, V. B. Shenoy, C. McCulloch, B. Hinz, Dynamic fibroblast contractions attract remote macrophages in fibrillar collagen matrix. Nat. Commun. 10 (2019), doi:10.1038/s41467-019-09709-6.

13. J. M. J. M. Szulczewski, D. R. D. R. Inman, D. Entenberg, S. M. S. M. Ponik, J. Aguirre-Ghiso, J. Castracane, J. Condeelis, K. W. K. W. Eliceiri, P. J. P. J. Keely, In Vivo Visualization of Stromal Macrophages via label-free FLIM-based metabolite imaging. Sci. Rep. 6, 25086 (2016).

14. J. S. Bredfeldt, Y. Liu, C. A. Pehlke, M. W. Conklin, J. M. Szulczewski, D. R. Inman, P. J. Keely, R. D. Nowak, T. R. Mackie, K. W. Eliceiri, Computational segmentation of collagen fibers from second-harmonic generation images of breast cancer. J. Biomed. Opt. 19, 016007 (2014).

15. S. P. Carey, Z. E. Goldblatt, K. E. Martin, B. Romero, R. M. Williams, C. A. Reinhart-King, Local extracellular matrix alignment directs cellular protrusion dynamics and migration through Rac1 and FAK. Integr. Biol. 8, 821–835 (2016).

16. D. Riveline, E. Zamir, N. Q. Balaban, U. S. Schwarz, T. Ishizaki, S. Narumiya, Z. Kam, B. Geiger, A. D. Bershadsky, contacts as mechanosensors: Externally applied local mechanical force induces growth of focal contacts by an mDia1-dependent and ROCK-independent mechanism. J. Cell Biol. (2001), doi:10.1083/jcb.153.6.1175.

17. N. Q. Balaban, U. S. Schwarz, D. Riveline, P. Goichberg, G. Tzur, I. Sabanay, D. Mahalu, S. Safran, A. Bershadsky, L. Addadi, B. Geiger, Force and focal adhesion assembly: A close relationship studied using elastic micropatterned substrates. Nat. Cell Biol. (2001), doi:10.1038/35074532.

18. B. A. Roeder, K. Kokini, J. E. Sturgis, J. P. Robinson, S. L. Voytik-Harbin, Tensile mechanical properties of three-dimensional type I collagen extracellular matrices with varied microstructure. J. Biomech. Eng. (2002), doi:10.1115/1.1449904.

19. S. Motte, L. J. Kaufman, Strain stiffening in collagen i networks. Biopolymers (2013), doi:10.1002/bip.22133.

20. O. V. Kim, R. I. Litvinov, J. W. Weisel, M. S. Alber, Structural basis for the nonlinear mechanics of fibrin networks under compression. Biomaterials (2014), doi:10.1016/j.biomaterials.2014.04.056.

21. A. J. Licup, S. Münster, A. Sharma, M. Sheinman, L. M. Jawerth, B. Fabry, D. A. Weitz, F. C. MacKintosh, Stress controls the mechanics of collagen networks. Proc. Natl. Acad. Sci. U. S. A. (2015), doi:10.1073/pnas.1504258112.

22. B. Burkel, M. Proestaki, S. Tyznik, J. Notbohm, Heterogeneity and nonaffinity of cell-induced matrix displacements. Phys. Rev. E. 98 (2018), doi:10.1103/PhysRevE.98.052410.

23. B. Burkel, J. Notbohm, Mechanical response of collagen networks to nonuniform microscale loads. Soft Matter. 13 (2017), doi:10.1039/c7sm00561j.

24. M. Proestaki, A. Ogren, B. Burkel, J. Notbohm, Modulus of Fibrous Collagen at the Length Scale of a Cell. Exp. Mech. 59, 1323–1334 (2019).

25. E. Bar-Kochba, J. Toyjanova, E. Andrews, K.-S. Kim, C. Franck, A Fast Iterative Digital Volume Correlation Algorithm for Large Deformations. Exp. Mech. 55, 261–274 (2015).

26. J. D. Eshelby, The Elastic Field Outside an Ellipsoidal Inclusion. Proceedings of the Royal Society of London, A252 (1271), 561–569. 252, 1959 (1959).

27. P. Rosakis, J. Notbohm, G. Ravichandran, A model for compression-weakening materials and the elastic fields due to contractile cells. J. Mech. Phys. Solids. 85, 16–32 (2015).

28. I. Acerbi, L. Cassereau, I. Dean, Q. Shi, A. Au, C. Park, Y. Y. Chen, J. Liphardt, E. S. Hwang, V. M. Weaver, and V. W. I Acerbi1, L Cassereau, I Dean, Q Shi, A Au, C Park, YY Chen, J Liphardt, ES Hwan, I. Acerbi, L. Cassereau, I. Dean, Q. Shi, A. Au, C. Park, Y. Y. Chen, J. Liphardt, E. S. Hwang, V. M. Weaver, Human breast cancer invasion and aggression correlates with ECM stiffening and immune cell infiltration. Integr. Biol. (United Kingdom). 7, 1120–1134 (2015).

29. K. Esbona, Y. Yi, S. Saha, M. Yu, R. R. Van Doorn, M. W. Conklin, D. S. Graham, K. B. Wisinski, S. M. Ponik, K. W. Eliceiri, L. G. Wilke, P. J. Keely, The Presence of Cyclooxygenase 2, Tumor-Associated Macrophages, and Collagen Alignment as Prognostic Markers for Invasive Breast Carcinoma Patients. Am. J. Pathol. (2018), doi:10.1016/j.ajpath.2017.10.025.

30. J. Wyckoff, W. Wang, E. Y. Lin, Y. Wang, F. Pixley, E. R. Stanley, T. Graf, J. W. Pollard, J. Segall, J. Condeelis, A paracrine loop between tumor cells and macrophages is required for tumor cell migration in mammary tumors. Cancer Res. (2004), doi:10.1158/0008-5472.CAN-04-1449.

31. S. Goswami, E. Sahai, J. B. Wyckoff, M. Cammer, D. Cox, F. J. Pixley, E. R. Stanley, J. E. Segall, J. S. Condeelis, Macrophages promote the invasion of breast carcinoma cells via a colony-stimulating factor-1/epidermal growth factor paracrine loop. Cancer Res. (2005), doi:10.1158/0008-5472.CAN-04-1853.

32. B. Burkel, J. Notbohm, Mechanical response of collagen networks to nonuniform microscale loads. Soft Matter. 13, 5749–5758 (2017).

33. D. Vader, A. Kabla, D. Weitz, L. Mahadevan, Strain-induced alignment in collagen gels. PLoS One. 4 (2009), doi:10.1371/journal.pone.0005902.

34. B. A. Roeder, K. Kokini, S. L. Voytik-Harbin, Fibril microstructure affects strain transmission within collagen extracellular matrices. J. Biomech. Eng. 131, 1–11 (2009).

35. M. D. C. Lopez-Garcia, D. J. Beebe, W. C. Crone, Mechanical interactions of mouse mammary gland cells with collagen in a Three-dimensional construct. Ann. Biomed. Eng. 38, 2485–2498 (2010).

36. H. Y. Yu, S. C. Sanday, C. I. Chang, Elastic inclusions and inhomogeneities in transversely isotropic solids. Proc. R. Soc. London. Ser. A Math. Phys. Sci. 444, 239–252 (1994).

