## Supplemental data for "Cell-Scale Biophysical Cues from Collagen Fiber Architecture Instruct Cell Behavior and the Propagation of Mechanosensory Signals"

### Supplementary Figures

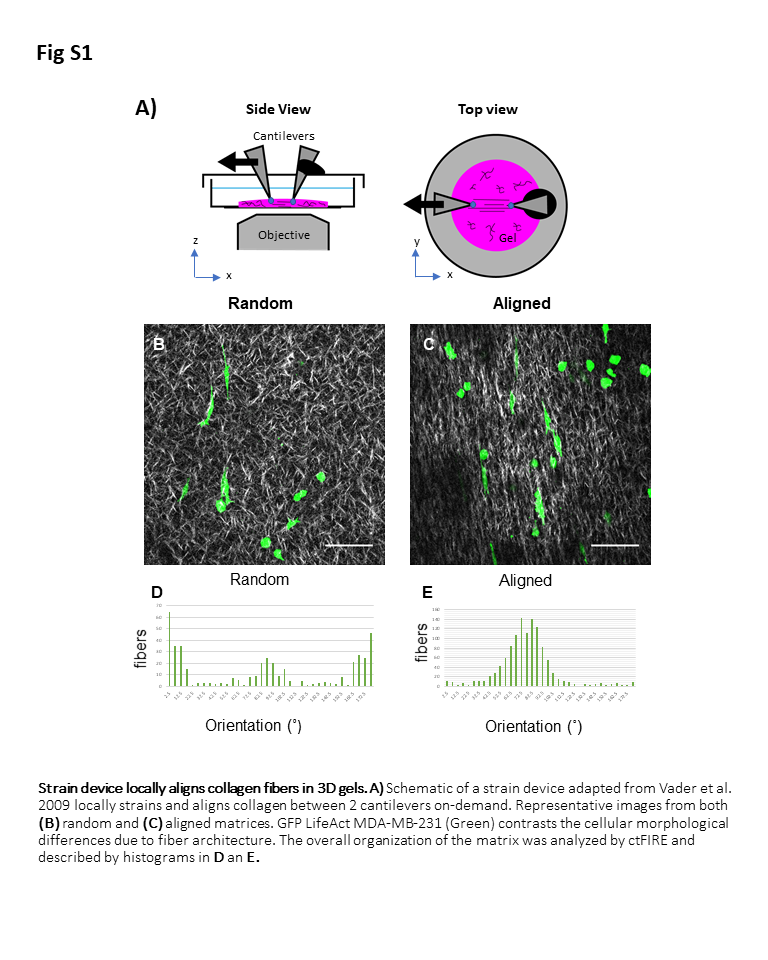

Figure S1: Strain device locally aligns collagen fibers in 3D gels. A) Schematic of a strain device adapted from Vader et al. 2009 (*33*) locally strains and aligns collagen between 2 cantilevers on-demand. Representative images from both (B) random and (C) aligned matrices. GFP LifeAct MDA-MB-231 (Green) contrasts the cellular morphological differences due to fiber architecture. The overall organization of the matrix was analyzed by ctFIRE and described by histograms in (D and E).

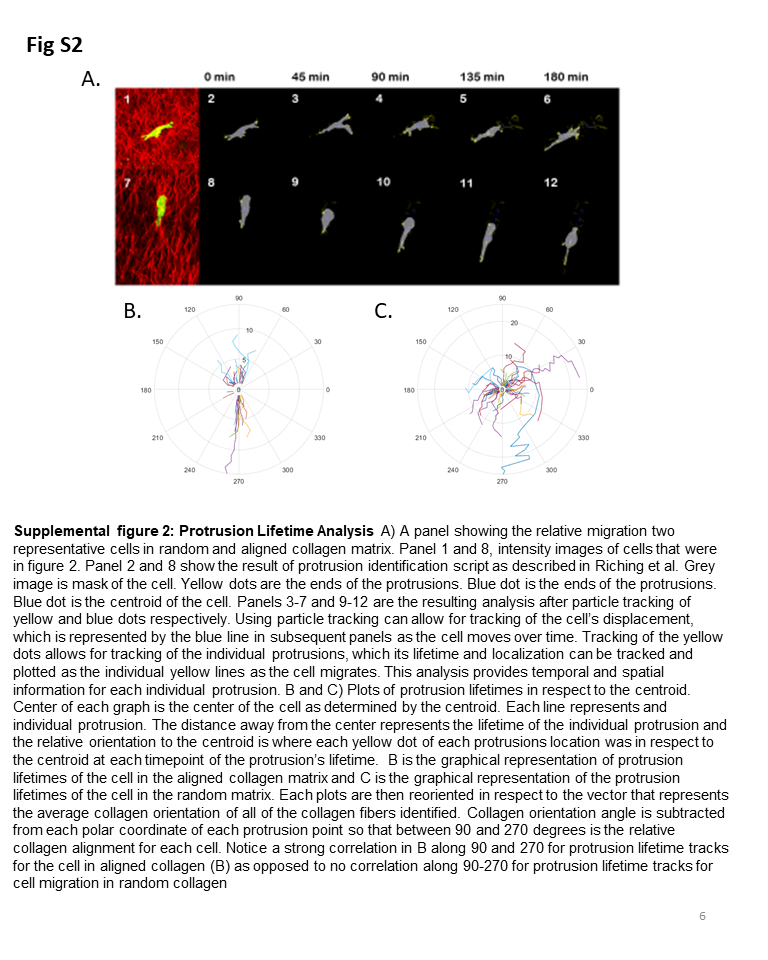

**Figure S2: Protrusion Lifetime Analysis** **A**) A panel showing the relative migration two representative cells in random and aligned collagen matrix. Panel 1 and 8, intensity images of cells that were in figure 2. Panel 2 and 8 show the result of protrusion identification script as described in Riching et al. Gray image is mask of the cell. Yellow dots are the ends of the protrusions. Blue dot is the ends of the protrusions. Blue dot is the centroid of the cell. Panels 3-7 and 9-12 are the resulting analysis after particle tracking of yellow and blue dots respectively. Using particle tracking can allow for tracking of the cell’s displacement, which is represented by the blue line in subsequent panels as the cell moves over time. Tracking of the yellow dots allows for tracking of the individual protrusions, which its lifetime and localization can be tracked and plotted as the individual yellow lines as the cell migrates. This analysis provides temporal and spatial information for each individual protrusion. **B and C)** Plots of protrusion lifetimes in respect to the centroid. Center of each graph is the center of the cell as determined by the centroid. Each line represents and individual protrusion. The distance away from the center represents the lifetime of the individual protrusion and the relative orientation to the centroid is where each yellow dot of each protrusions location was in respect to the centroid at each timepoint of the protrusion’s lifetime. **B)** graphical representation of protrusion lifetimes of the cell in the aligned collagen matrix and **C)** graphical representation of the protrusion lifetimes of the cell in the random matrix. Each plots are then reoriented in respect to the vector that represents the average collagen orientation of all of the collagen fibers identified. Collagen orientation angle is subtracted from each polar coordinate of each protrusion point so that between 90 and 270 degrees is the relative collagen alignment for each cell. Notice a strong correlation in B along 90 and 270 for protrusion lifetime tracks for the cell in aligned collagen (B) as opposed to no correlation along 90-270 for protrusion lifetime tracks for cell migration in random collagen

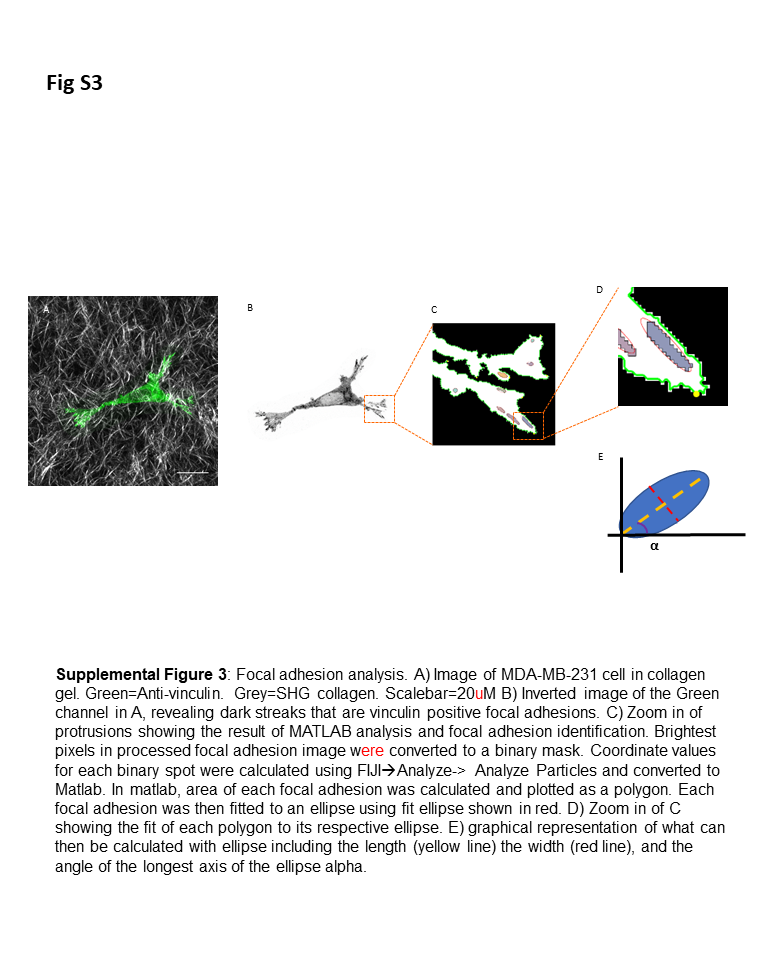

**Figure S3**: **Focal adhesion analysis.** **A)** Image of MDA-MB-231 cell in collagen gel. Green = Anti-vinculin. Gray = SHG collagen. Scalebar = 20 µM **B)** Inverted image of the Green channel in A, revealing dark streaks that are vinculin positive focal adhesions. **C)** Zoom in of protrusions showing the result of MATLAB analysis and focal adhesion identification. Brightest pixels in processed focal adhesion image were converted to a binary mask. Coordinate values for each binary spot were calculated using FIJI🡪Analyze-> Analyze Particles and converted to Matlab. In Matlab, the area of each focal adhesion was calculated and plotted as a polygon. Each focal adhesion was then fitted to an ellipse using fit ellipse shown in red. **D)** Zoom in of C showing the fit of each polygon to its respective ellipse. **E)** graphical representation of what can then be calculated with ellipse including the length (yellow line) the width (red line), and the angle of the longest axis of the ellipse alpha.

### Supplemental Note: Method to Compute Anisotropic Moduli

The contractile strains of the particle in directions parallel and perpendicular to fiber alignment were used to determine the moduli of the matrix in those two directions. Our method extended our previous work (*24*) to apply to an anisotropic material. The anisotropic material model chosen was that of transverse isotropy, which is isotropic in one plane and anisotropic out of that plane, accounting for the increase in stiffness due to fiber alignment. Here we express components of stress ***σ*** and strain ***ε*** in Voight notation wherein ***σ*** and ***ε*** are vectors containing the 6 independent components of stress and strain. Stress and strain are related through the symmetric 6 *×* 6 stiffness matrix ***C*** according to ***σ*** = ***Cε***.

We define unit vectors in the three Cartesian directions to be ***e***_1_, ***e***_2_, and ***e***_3_ and the direction of fiber alignment to be the ***e***_3_ direction. Then *C*_11_ = *C*_22_, *C*_13_ = *C*_23_, and *C*_44_ = *C*_55_ due to the in-plane isotropy. Additionally, due to the transverse anisotropy, *C*_66_ = (*C*_11_ – *C*_12_)*/*2. The main parameters of interest are the stiffness in directions perpendicular to and parallel to fiber alignment, *C*_11_ and *C*_33_, respectively. Other unknown constants are *C*_12_, *C*_13_, and *C*_44_. As *C*_12_ and *C*_13_ couple normal stresses and strains between different directions, they are related to the in-plane and out-of-plane Poisson’s ratios, *ν*_1_ and *ν*_31_ respectively. Here we used *ν*_1_ = 0*.*3 and *ν*_31_ = 2, which is consistent with previous experimental studies (*34*, *35*). Following standard equations of transverse isotropy, we related *C*_12_ and *C*_13_ to *C*_11_ and *C*_33_ (Fig. S4). Final values used in the analysis were *C*_12_ = 0*.*3 Pa and *C*_13_ = 2 Pa, which are consistent with the final computed elastic constants *C*_11_ and *C*_33_ (Fig. S4). The remaining parameter *C*_44_ was unknown, so we chose a range of different values for *C*_44_ and verified that the general trends were insensitive to the specific value.

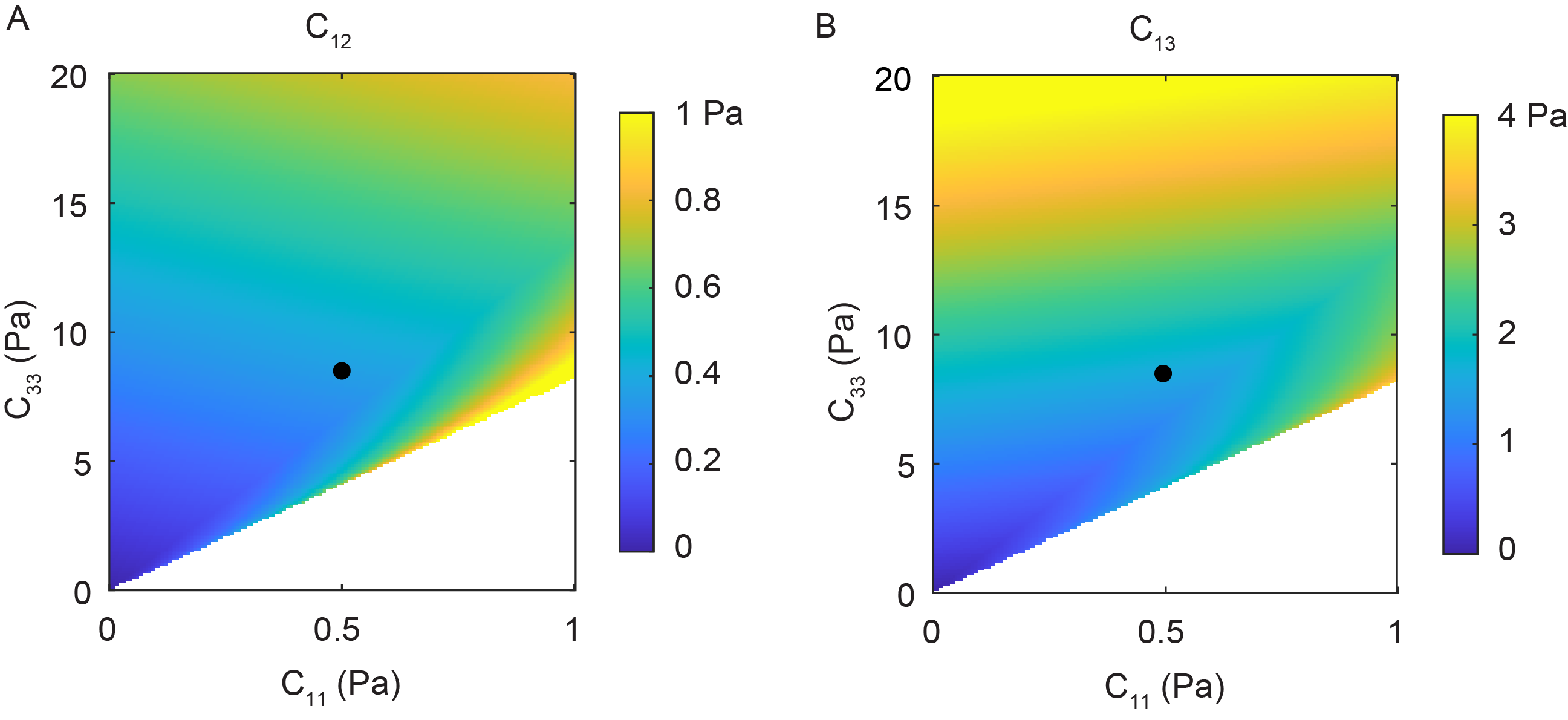

**Figure S4: Effect of moduli *C*_11_ and *C*_33_ on *C*_12_ (A) and *C*_13_ (B).** These graphs use values of in-plane and out-of-plane Poisson’s ratios 0.3 and 2, respectively. Black dots show values of *C*_12_ and *C*_13_ chosen for the analysis, which correspond to *C*_11_ ≈ 0.5 and *C*_33_ ≈ 8 Pa, which are consistent with the experimentally measured values in Fig. 4C.

To compute *C*_11_ and *C*_33_ from strains of the particle, we used the linear elasticity solution for a sphere contracting in a transversely isotropic medium (*36*). As the equations relating contractile strain to matrix moduli are nonlinear, it is unknown whether a closed form solution exists to compute moduli from particle strains. Therefore, we solved iteratively using the following approach:

1. Elastic moduli *C*_11_ and *C*_33_ were chosen.
2. The Eshelby tensors were computed using expressions reported by Refs. (*36*) and (*26*).
3. Note that the PNIPAAm particle is isotropic and therefore has different elastic constants than the surrounding matrix, whereas the classical Eshelby solution assumes the contracting inclusion and matrix to have the same moduli. Therefore, the Eshelby tensors were combined with the measured thermal contraction and elastic moduli of the particle to compute the equivalent transformation strain for a contracting inclusion having the same elastic properties as those of the matrix as described by Ref. (*36*).
4. The contractile strains of the inclusion in directions parallel and perpendicular to fiber alignment were computed and compared to the measured values.
5. The process was repeated for different moduli *C*_11_ and *C*_33_. The values of *C*_11_ and *C*_33_ that minimized the difference between computed and measured strains were taken to be the measured values.

One additional consideration is that fibrous materials are nonlinear, with elastic moduli far lower in compression compared to tension. Using the solution for a compression softening material (*27*), we showed in our previous work (*24*) that for an isotropic matrix, the effect of compression softening is equivalent to in- creasing the bulk modulus of the particle by a factor of 2. As there is no solution for a sphere contracting in a compression softening anisotropic medium, we reasoned that to first order, compression softening would have the same effect in isotropic and anisotropic matrices. Hence, in computing the moduli of the matrix, we multiplied the bulk modulus of the PNIPAAm particles by a factor of 2.

As in our previous study (*24*), we accounted for experimental uncertainty due to particle-to-particle variability in particle moduli and thermal contraction using a bootstrap analysis. The analysis gave estimates of the mean along with 95% confidence intervals of the mean of modulus of matrix in the local region surrounding each particle.

Fig. S5 shows results for three different values of shear modulus *C*_44_: 2, 4, and 8 Pa. For all values of *C*_44_, trends are qualitatively and quantitatively the same—the modulus parallel to fiber alignment *C*_33_ ranges from 5 to 12 Pa, and the ratio *C*_33_*/C*_11_ is linearly proportional to *C*_33_. Hence, the results are insensitive to the choice of *C*_44_; the central value, *C*_44_ = 4 Pa was chosen to present in the main text.

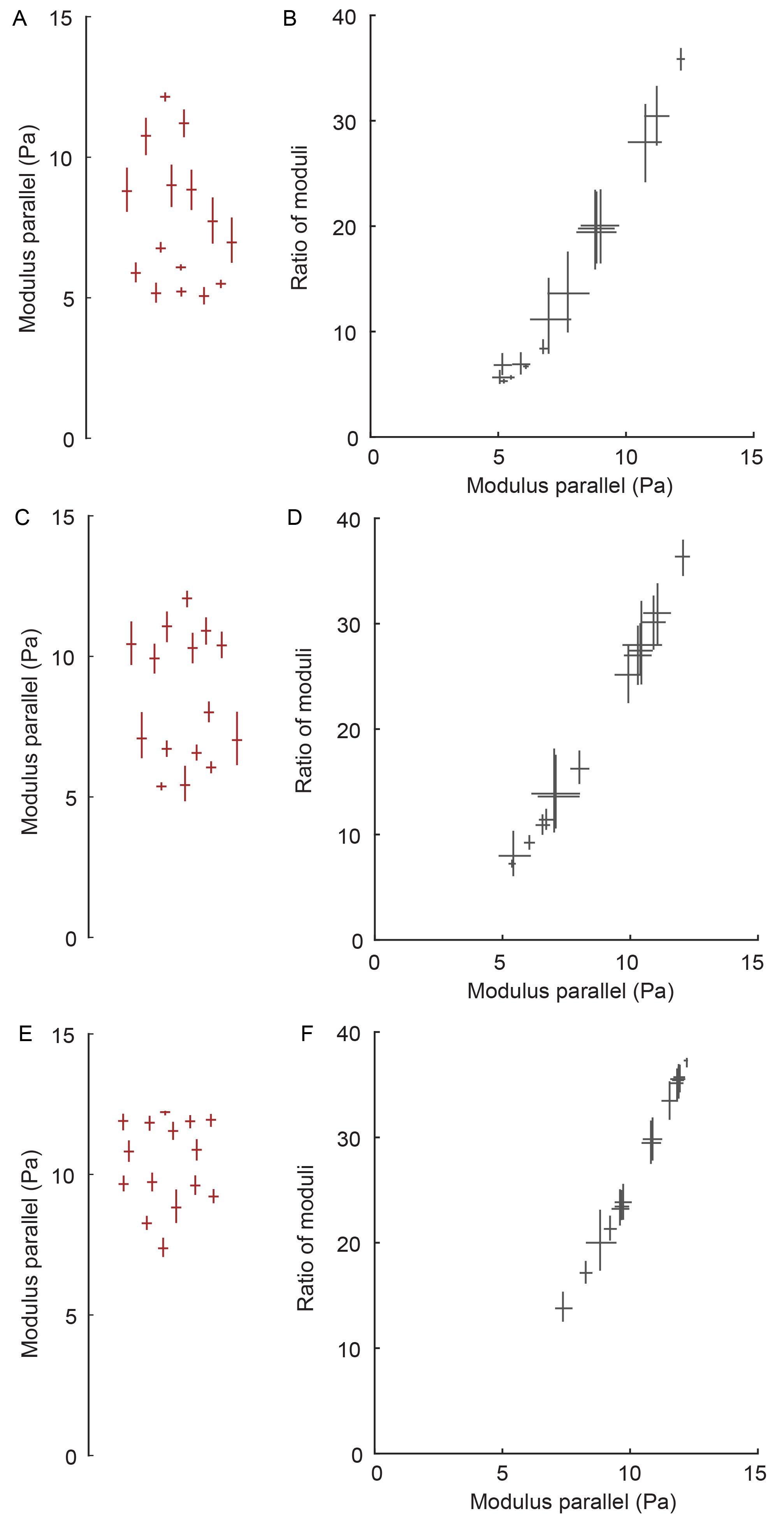

**Figure S5: Effect of shear modulus *C*_44_.** Because the shear modulus *C*_44_ is unknown, the moduli parallel and perpendicular to fiber alignment were computed for three different values of *C*_44_: 2 Pa **(A, B)**, 4 Pa **(C, D)**, and 8 Pa **(E, F)**. In all cases, trends in the data remain the same, indicating that the results are insensitive to *C*_44_. The central value, *C*_44_ = 4 Pa was chosen to present in the main text, so panels C and D are the same as those in Fig. 4.
